# Identification of Inhibitors of *Chikungunya* virus nsP2 ATPase

**DOI:** 10.1101/2024.12.02.625520

**Authors:** Hernan Navarro, John E. Scott, Ginger R. Smith, Pegah Ghiabi, Elisa Gibson, Peter Loppnau, Rachel J. Harding, Mohammad Anwar Hossain, Muthu Ramalingam Bose, Kenneth H. Pearce, Eric M. Merten, Timothy M. Willson, Peter J. Brown

## Abstract

Non-structural protein 2 (nsP2), which plays an essential role in replication of CHIKV, contains a protease, helicase, and methyltransferase-like domain. We executed a simple a screen using malachite green to detect compounds that decreased ATP hydrolysis and tested a library of diverse compounds to find inhibitors of CHIKV nsP2 helicase.

## Introduction

*Chikungunya* virus (CHIKV) infections have spread among the Americas, Africa and Asia by infected mosquitos resulting in fever and joint pain and swelling. Non-structural protein 2 (nsP2), which plays an essential role in replication of CHIKV, contains a protease, helicase, and methyltransferase-like domain. The helicase domain is a motor enzyme that consumes ATP as a cofactor as it unwinds the RNA duplex during viral genome replication. As viral helicases are involved in the replication mechanism of viruses, inhibition of viral helicases is a viable strategy for developing antiviral agents. A few inhibitors are known for human helicases^1-11^, whereas none has been published for viral helicases. Nsp2 protease is responsible for cleaving viral polyprotein into the competent enzymes of the replication process (nsp1,2,3,4)^12^. To identify potential nsP2 helicase inhibitors, we executed a simple a screen using malachite green^13^ to detect compounds that decreased ATP hydrolysis. A screen of a 48,712 member library from ChemDiv identified several potential nsP2 helicase inhibitors.

### Primary Assay

An ATPase assay was developed using Malachite Green as a detector of free phosphate and a collection of 48,712 diverse compounds from ChemDiv was screened at 25 μM. The Malachite Green assay performed with an average plate-based Z’-factor of 0.7. A cut-off of ≥46% inhibition was applied to identify plates with initial actives. Entire compound plates with any active compounds were then re-screened to confirm actives. After this replicate screen, hits were tested in a dose-response manner in a secondary assay employing ADP-Glo.

### Secondary Assay

A CHIKV nsp2 ATPase assay was developed using ADP-Glo and Kinase-Glo reagents from Promega.

## Results

30 confirmed actives were identified from the primary assay with repeat activity at 25 μM for a confirmed hit rate of 0.06%. These compounds were resupplied for testing in the secondary assay (Table 1). Of the 30 compounds followed up by dose response in the ADP-Glo assay, nine showed IC_50_ < 10 μM with acceptable Hill slopes (0.5-2.0) and one with Hill slope 2.8. The most potent compound, RA-0001819 (IC_50_ 0.6 μM) was a highly substituted sulfonamide which was not considered tractable from a MedChem perspective with medium kinetic solubility of 12.5 μM. The next most potent template showed up three times in the top six compounds, RA-0001821, 1822 and 1823 with IC_50_s of 4.0, 2.3 and 1.5 μM respectively. These compounds showed medium to high kinetic solubility (14.2, 236 and 202 μM respectively). Several compounds containing an alkylidene barbiturate moiety also inhibited ATPase activity with IC_50_s below 10 μM (RA-0001799, 1798 and 1800), however, these are considered Pan Assay Interference compounds (PAINS)^14^ and were not investigated further.

**Table 1.**
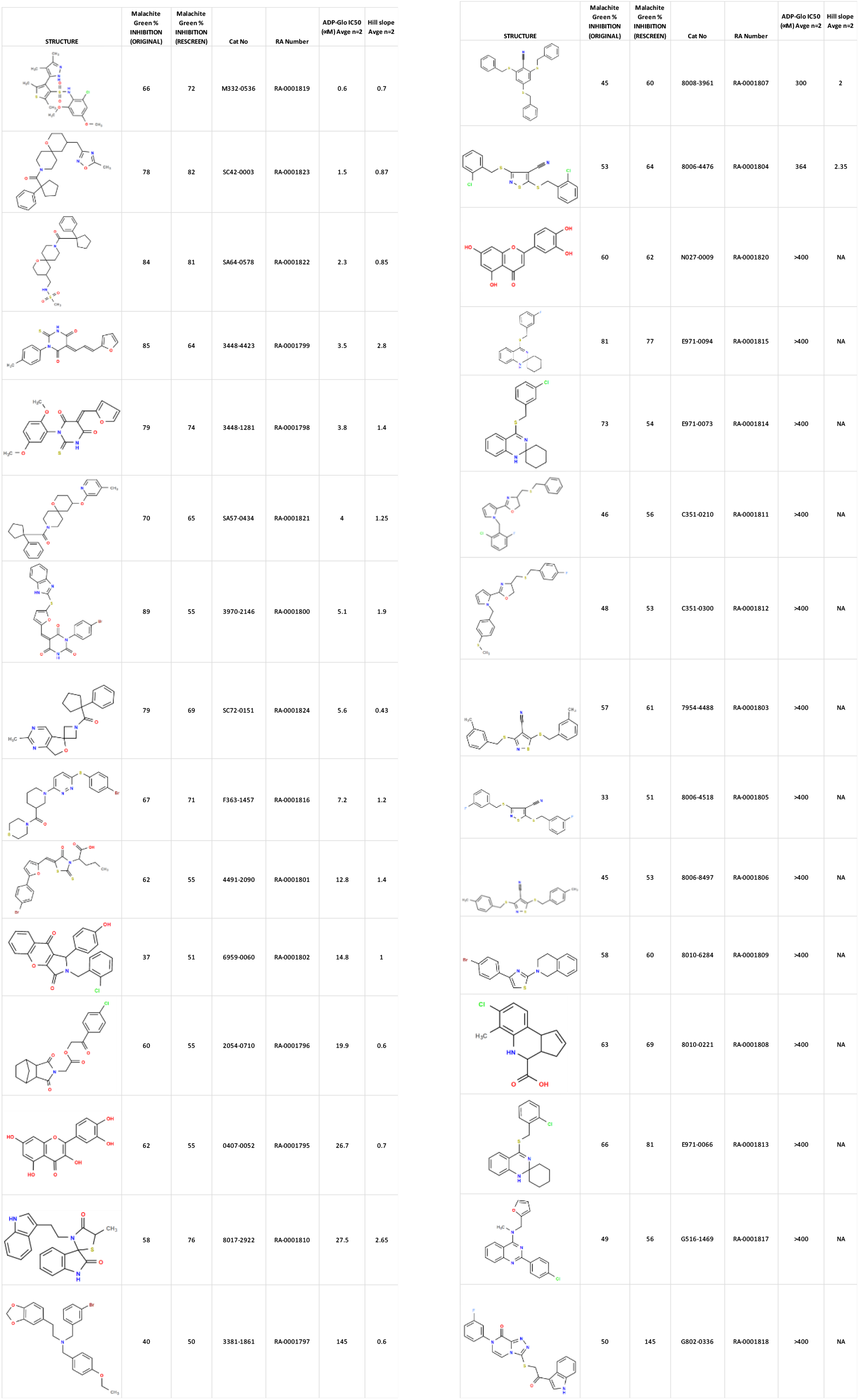
Structures and related activity data from malachite green HTS and ADP-Glo follow-up assays.

## Conclusion

Using a simple assay for inorganic phosphate production, several inhibitors of CHIKV nsP2 ATPase activity were identified. A spiropiperidine chemotype was identified multiple times among the most potent of the ATPase inhibitors. Additional studies will be required to demonstrate the mechanism of action of these spiropiperidines as potential inhibitors of nsP2 RNA helicase.

## Materials and Methods

### CHIKV nsp2 Protein Expression and Purification

cDNA corresponding to aa. 536-1333 of the CHIKV polyprotein was subcloned into a modified version of the pET28-MHL vector that yields a N-terminal TEV cleavable his-tag and C-terminal thrombin cleavable avi-tag. The protein was coexpressed with BirA in *E. coli* BL21 codonplus (DE3), grown in TB, to produce biotinylated protein. Cell pellets were re-suspended in purification buffer 1 (20 mM HEPES pH 6.8, 500 mM NaCl, 20% (v/v) glycerol, 2 mM MgCl_2_, 1 mM TCEP, 5 mM imidazole) and supplemented with 0.5% (v/v) CHAPS, 1 mM PMSF/Benzamidine and 100 µL 0.1 mg/mL benzonase (produced in-house) and lysed by sonication. The crude extract was clarified by high-speed centrifugation (60 min at 36,000 ×g at 4 °C) and the clarified lysate was batch adsorbed, while rotating at 4 °C, with pre-equilibrated Ni-NTA regenerated for 30 mins. The resin was washed with purification buffer 1 supplemented with 1 mM biotin, followed by another wash using purification buffer 1 supplemented with 20 mM imidazole. Finally, the protein was eluted by purification buffer 1 supplemented with 250 mM imidazole. The eluted protein was applied to a HiLoad Superdex200 26/600 using an ÄKTA Pure (Cytiva) pre-equilibrated with purification buffer 2 (50 mM HEPES pH 7, 150 mM NaCl, 10% (v/v) glycerol, 2 mM MgCl_2_, 1mM TCEP). Fractions containing nsp2 protein solution were filtered and loaded onto an 8 mL CaptoHiRes S cation exchange column. The column was washed with 15% B over 5 CV and then eluted over 20 CV up to 50% B buffer. (Buffer A: purification buffer 2 with 0 mM NaCl / Buffer B: purification buffer 2 with 1M NaCl). The peaks eluting from the column at ∼300 mM NaCl were pooled and concentrated. The concentration was then measured by nanodrop, the protein was aliquoted and then flash frozen in liquid nitrogen for storage at -80 °C.

### High Throughput Screen with the nsP2 ATPase Activity Assay (Malachite Green)

The high throughput nsP2 ATPase activity assay was performed in 384-well plates (Corning white walled, clear bottom cat # 3763). A Nanoscreen NSX 1536 (Charleston, SC) liquid handling instrument equipped with a 384-tip head was used to transfer 10 μL of assay buffer (25 mM HEPES pH 7.5, 5 mM MgCl_2_, 1 mM DTT) to each well. Subsequently, 5 μL of purified nsP2 enzyme diluted into assay buffer was added to each well. 50 nL of library compound in 100% DMSO (or 100% DMSO for control wells) was added to each well using a Biomek NX (Beckman Coulter Inc., Fullerton, CA) equipped with a pin tool head (V&P Scientific, San Diego, CA). The plates were placed on a plate shaker for 1 minute. The enzyme reaction was initiated by the addition of 5 μL of ATP diluted into assay buffer resulting in final concentrations of 1 mM ATP and 3 nM enzyme in a final volume of 20 μL. Final concentration of compound was 25 μM and DMSO was 0.25% in all wells. Plates were then put on a plate shaker for 1 minute and incubated at room temperature for 30 minutes. Subsequently, 40 μL of malachite green reagent (Bioassay Systems kit, Hayward, CA) was transferred to each well to detect free phosphate. Plates were incubated in the dark for 30 minutes and absorbance (620 nm) measured on a SpectraMax 384 Plus plate reader. Absorbance values were normalized to the mean of DMSO (100% activity) and “no enzyme” (0% activity) control wells as maximum and minimum signals, respectively, to obtain percent inhibition values. Z’-factor calculations based on plate controls were performed.

#### Secondary CHIKV nsP2 ATPase assay

The ADP-Glo Kinase assay kit (Cat# V9101) was purchased from Promega and used to evaluate compounds for inhibition of ATPase activity. Active compounds from the ChemDiv library screen were repurchased from ChemDiv. Enzymatic reactions were performed in 384-well, white, low volume ProxiPlates (Revvity, Cat# 6008280). To prepare dose-response plates, ten-point, three-fold compound dilutions were prepared in DMSO using a Tecan Evo, beginning from a top concentration of 10 mM. Using a Mosquito liquid handler (STP LabTech), 200 nL of each dilution were transferred to assay-ready plates. Assays were performed with a final volume of 10 µL, resulting in a final top concentration of 200 µM. For the assay, a stock solution of 5X activity buffer (200 mM Tris pH 7.5, 0.5 mg/mL BSA) was prepared and filtered through a 0.2 μm filter. Assay buffer was prepared from 100 mM MgCl_2_, 1M DTT, and the 5x activity buffer stock, for a final 1x assay buffer composition of 40 mM Tris pH 7.5, 0.1 mg/mL BSA, 2 mM MgCl_2_, and 1 mM DTT. A 2x solution of CHIKV nsP2fl (2 nM, 1 nM final) was prepared in 1x assay buffer. 5 µL of 2x CHIKV nsP2fl was added to pre-plated compounds and pre-incubated for 30 minutes at room temperature. A 2x solution of Ultra-Pure ATP (60 µM, 30 µM final) was prepared in 1x assay buffer. After the 30-minute pre-incubation with enzyme, 5 µL of 2x ATP was dispensed into the assay plate to initiate the enzymatic reaction. Reagents were dispensed into assay plates using a Combi Multidrop (Thermo Scientific). The reactions were allowed to proceed for 1 hour, and then the reactions were quenched by the addition of 2 µL of ADP-Glo reagent to deplete remaining ATP for 40 minutes. Next, 2 µL of Kinase Glo reagent was added and incubated for 30 minutes. Luminescence values were determined using a PerkinElmer Envision 2105 plate reader. Percent inhibition values and Z’ scores were calculated using a no-enzyme column and a DMSO-only column as low and high controls, respectively.

